# NicheTrans: Spatial-aware Cross-omics Translation

**DOI:** 10.1101/2024.12.05.626986

**Authors:** Zhikang Wang, Senlin Lin, Qi Zou, Yan Cui, Chuangyi Han, Yida Li, Jianmin Li, Yi Zhao, Rui Gao, Jiangning Song, Zhiyuan Yuan

## Abstract

Spatial omics technologies have revolutionized our studies on tissue architecture and cellular interactions at single-cell resolution. While spatial multi-omics approaches offer unprecedented insights into complex biological systems, their widespread adoption is hindered by technical challenges, specialized requirements, and limited accessibility. To address these limitations, we present NicheTrans, the first spatially-aware cross-omics translation method and a flexible Transformer-based multi-modal deep learning framework. Unlike existing single-cell (non-spatial) translation methods, NicheTrans uniquely incorporates both cellular microenvironment information and flexible integration of multi-modal data, such as morphology and prior knowledge. We validated NicheTrans across diverse biological cases: Parkinson’s Disease (PD), Alzheimer’s Disease (AD), breast cancer, and lymph nodes. Our approach demonstrated superior performance compared to existing single-cell methods, highlighting the crucial role of spatial and multi-modal information in cross-omics translation. Through NicheTrans, we uncovered spatial multi-omics domains that were not detectable through single-omics analysis alone. Model interpretation revealed key molecular relationships, including gene programs associated with dopamine metabolism and amyloid β-associated cell states. Additionally, using translated protein markers as spatial landmarks, we quantified the spatial organization of key glial cell subtypes in the AD brain. NicheTrans represents a powerful tool for generating comprehensive spatial multi-omics insights from more accessible single-omics measurements, making multi-omics analysis more feasible for the broader research community.

## 1. INTRODUCTION

The spatial organization of cells within tissues provides essential insights into cellular interactions, tissue architecture, and disease mechanisms^1–3^. Advances in spatial omics have enabled the spatial measurement of gene expression, proteins, and/or metabolic features at up to single-cell resolution, leading to novel insights across fields including neuroscience, developmental biology, disease, and cancer^4–6^. Notably, spatially resolved transcriptomics has undergone significant development in recent years^7^, improving in spatial resolution^8,9^, gene expression coverage^10^, field of view size^9,11^, three-dimensional capabilities^12–15^, and experimental feasibility^16,17^. Datasets resolving specific omics layers are now readily available through large consortia^1,18–21^.

However, the complexity of biological systems demands more than single-omics information alone^22^. This recognition has driven the development of spatial multi-omics technologies, which can simultaneously profile multiple molecular layers within the same tissue. Notable techniques have been developed for spatially co-mapping protein and gene expression^23,24^, metabolite and gene expression^25^, as well as chromatin accessibility and gene expression^26^. These technologies have provided unprecedented opportunities for characterizing and investigating the multi-omics molecular landscape of complex tissues with spatial resolution^27^.

Despite the development of these spatial multi-omics techniques, their widespread adoption faces significant challenges. First, their successful implementation depends on meticulous preparation under highly controlled experimental conditions, making them prone to technical variations. For example, the spatial multimodal analysis (SMA) protocol requires three washes in cold methanol at the end of mass spectrometry imaging (MSI) by matrix-assisted laser desorption/ionization^25^. This step can cause tissue degradation, reducing the availability of RNA molecules for sequencing. Second, spatial multi-omics experiments are more technically challenging than spatial single-omics ones, involving trade-offs between resolution, sensitivity, and throughput. Third, many of these technologies remain confined to a few specialized top laboratories and have not been commercialized, limiting their availability and accessibility to the broader research community.

The single-cell (non-spatial) omics field has addressed similar multi-omics challenges through computational approaches^28^. Methods like cTP-net^29^, sciPENN^30^, and JAMIE^31^ have successfully demonstrated cross-omics translation in non-spatial contexts, using deep learning to predict protein abundances from RNA data or to integrate chromatin accessibility information (see Supplementary Note 1 for a more detailed review). However, applying these methods directly to spatial data is suboptimal for several reasons. First, cellular phenotypes in tissue contexts are influenced not just by intrinsic gene expression but also by the surrounding microenvironment – cells with similar transcriptional profiles can exhibit different protein or metabolic signatures based on their spatial context (e.g., proximity to blood vessels)^32^. Second, spatial technologies often generate additional data modalities, such as histological images, which could provide critical information for cross-omics translation but are not utilized by existing methods^25^. There is a pressing need for a spatial-aware cross-omics translation method that can integrate both cell niche information and additional data modalities.

Here we developed NicheTrans, a Transformer-based multi-modal deep learning framework specifically designed for spatial multi-omics translation. In the computation process, spatial information is incorporated by formulating the spatial context of each cell as a niche. Multi-modal data, including measured omics data, cell type labels (if available), and Hematoxylin and Eosin (H&E) images (if available), can be flexibly integrated as input features. We validated and benchmarked NicheTrans using four diverse spatial multi-omics datasets spanning Parkinson’s Disease (PD), Alzheimer’s Disease (AD), breast cancer, and lymph node tissues. The results demonstrated superior performance across various evaluation metrics compared to existing methods. The utility of NicheTrans extends beyond simple cross-omics translation. Using our translation, we identified spatial domains characterized by integrated gene expression and metabolic signatures that were obscured in single-omics analysis alone. We utilized the model interpretation method to reveal key relationships between different molecular layers, such as gene programs associated with dopamine metabolism and amyloid β (Aβ)-associated cell states. Furthermore, using translated protein markers as spatial landmarks, we quantified the spatial organization of key subtypes of glial cells in the AD brain. We additionally evaluated the superior performance in non-stereotypical tissues other than the brain. In all, NicheTrans facilitates the generation of comprehensive spatial multi-omics data from more readily available single-omics measurements, making multi-omics insights more accessible to the broader research community.

## 2. RESULTS

### Motivation

We explore alternatives to achieve spatial-aware cross-omics translation. The first approach involves applying single-cell non-spatial translation methods^29–31^ to spatial data, without considering spatial information. However, this approach has limitations because cellular states within tissues are significantly influenced by their microenvironment, indicating that cells with similar gene expressions but different locations may exhibit different phenotypes such as protein or metabolic states^33,34^. In Figure. 1A, we display cells (excluding those with zero expression) that have identical measured gene expression profiles across the Merscope (Vizgen), Xenium (10x)^35^, and Enhanced ELectric Fluorescence in situ Hybridization (EEL FISH)^36^ platforms (see Methods). Each point represents a distinct gene expression profile (the y-axis represents the number of cells that share the profile). Figure 1B shows the spatial distribution of gene expression-identical cells in a Merscope (Vizgen) human ovarian cancer sample (see Methods). Different colors represent unique gene expression profiles, and spot sizes indicate the number of cells sharing these profiles. Importantly, these identical cells are distributed across various locations within the tissue (Figure. 1B). This may contribute to different cellular phenotypes in terms of protein or metabolic state. However, using single-cell translation methods that only consider omics information (e.g., the measured gene expression) as input, without accounting for spatial information, will result in identical translation output inaccurately.

**Figure. 1:**
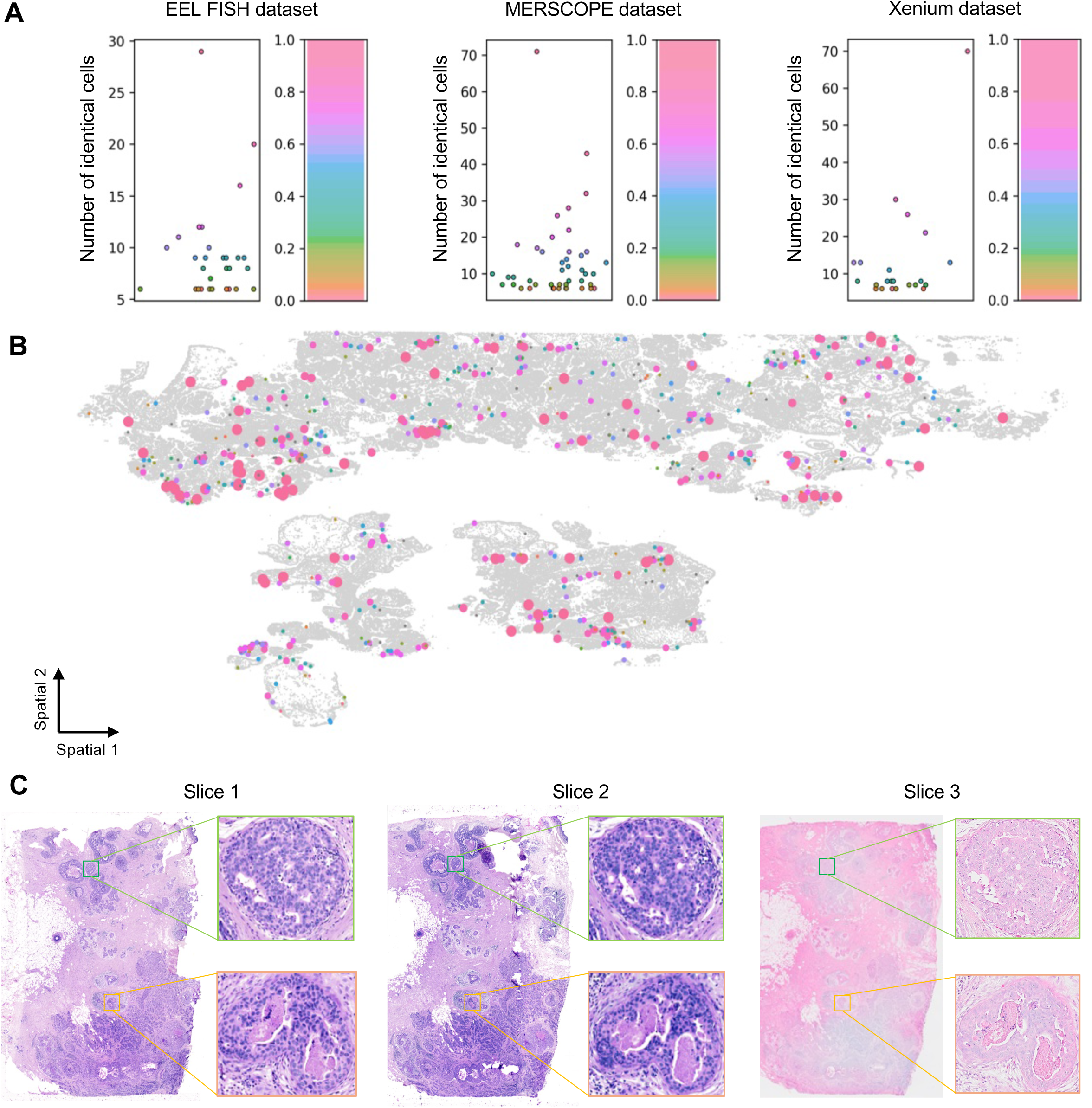
Motivation for spatial multi-omics translation methods. A. Dot plots showcasing cells with identical gene expression patterns across three image-based sequencing datasets. Each dot represents a specific gene expression pattern, with the y-axis indicating the number of cells sharing that pattern. B. Visualization of ovarian cancer tissue from the MERSCOPE dataset. Spot colors correspond to the dot plot above, while spot sizes indicate the number of cells with the associated gene expression pattern. C. Visualization of three serial slices from the Xenium breast cancer dataset, including zoomed-in views of two selected regions. The insets reveal micro-scale differences in cellular quantity, density, and spatial distribution.

Another alternative approach for spatial multi-omics involves computationally aligning two adjacent slices analyzed with two different spatial assays^37,38^. For example, one slice undergoes spatial transcriptomics while the adjacent one undergoes spatial proteomics (as demonstrated in two recent studies^2,39^). These slices are then spatially aligned to create two-omics data for each cell by identifying its nearest pair to complement the missing omics information. However, existing spatial alignment methods are typically designed for the same omics type, and methods for aligning different spatial omics are lacking^40^. Furthermore, while adjacent slices show structural similarity at the macroscopic level, their micro-level cellular distributions can differ significantly. Adjacent slices also inevitably experience shifts that make cell-to-cell matching impossible^39^. In Figure 1C, we present three serial sections of breast cancer tissue with 5 μm intervals and particularly zoom in two selected regions. While the tissue structures at the macroscopic level appear highly similar between slices, cell-to-cell matching at the microscopic level is not feasible due to the difference in local cellular quantity, density, and distribution (Figure. 1C). This challenge is exacerbated when slices with complementary spatial assays exhibit greater structural differences.

### NicheTrans overview

NicheTrans is a Transformer-based multi-modal deep neural network designed to translate between different types of omics data with spatial resolution, generating spatial multi-omics data (Figure. 2). For example, translating gene expression data into proteins or metabolites (Figure. 2C). The spatially resolved dependency information within each microenvironment is automatically learned and particularly emphasized to enable precise cross-omics translation. Because of its versatile design, NicheTrans can handle different spatial omics data, so “spot” (referring to larger tissue areas in low-resolution technologies)^41–43^ and “cell” (referring to individual cells in high-resolution technologies)^12,19,44^ are used interchangeably in this manuscript.

**Figure. 2:**
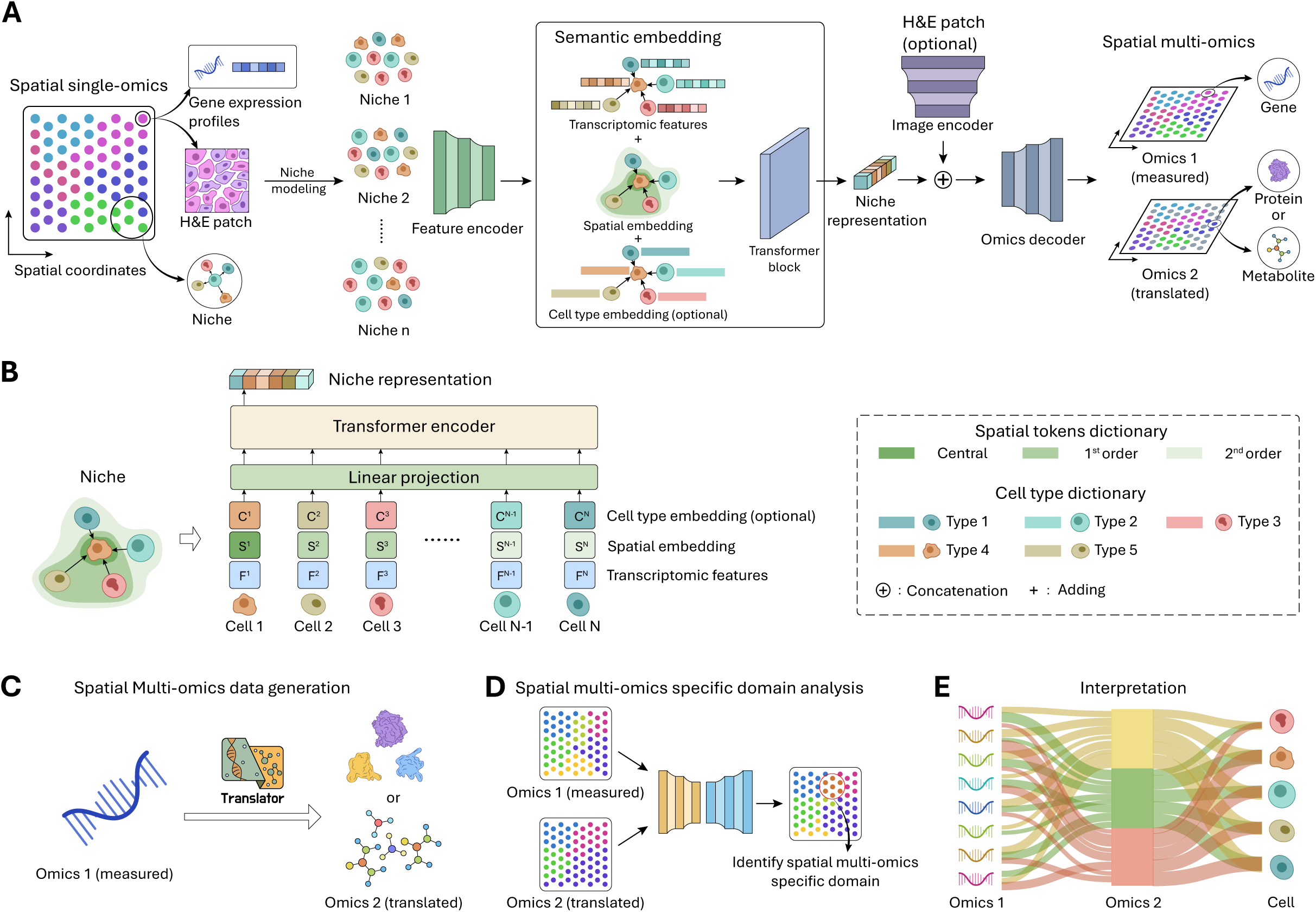
Schematic overview of NicheTrans and its downstream applications. A. Workflow of NicheTrans: NicheTrans translates source omics (e.g., gene expression) into target omics (e.g., proteomics or metabolomics) with spatial context. The input to NicheTrans comprises spatial single-omics data, with optional integration of cell type labels, spatial context, and morphological features from H&E images to enhance translation performance. B. The semantic embedding operation encodes spatial context and cell type information within each niche, while the Transformer encoder facilitates cell-cell communication throughout the translation process. C. NicheTrans enables the generation of spatial multi-omics data through cross-omics translation. D. Spatial multi-omics data generated by NicheTrans reveals specific spatial domains obscured in single-omics analysis alone. E. Model interpretation reveals correlations between omics layers and between omics and cell types.

Built on the Transformer architecture^45^, NicheTrans excels at cell-level feature learning and modeling complex spatial organizations. As shown in Figure 2A, the spatial single-omics data (a tissue slice assayed by a spatial single-omics technology) was first transformed into niches, where each niche comprises a center cell and its spatially adjacent neighboring cells. For illustration purposes, we used spatial transcriptomics data (spatial gene expression measurements for each cell) as an input example. During forward computation, NicheTrans employs a feature encoder to transform the input data into feature vectors, which focus on modeling gene-gene dependencies during translation. Subsequently, the semantic embedding operation (Figure. 2B) integrates different types of information into a unified representation, combining transcriptomic information, spatial information, cell type information (if provided, optional), and any other relevant prior knowledge associated with each niche. A Transformer block, comprising a multi-head self-attention module and a feed-forward network, is then used to capture the multi-modal dependencies within each niche (Figure. 2A). The features associated with the central cell are utilized as the final “niche representation” (Figure. 2A). Additionally, morphological features extracted from H&E images can be incorporated, if available (e.g., 10x Visium, Xenium, etc.), to provide additional information for translation (Figure. 2A). These morphological features are processed using convolutional neural networks (CNNs)^46^. Finally, the omics decoder that comprises multiple subnetworks will translate the input transcriptomics data to the target omics, generating the spatial multi-omics data (Figure. 2A).

During the training phase, spatial multi-omics data is used to optimize the neural network by designating one omics data as input and another as the target labels. In the testing phase, NicheTrans translates the input omics to the target omics, enabling in situ multi-omics analysis. For detailed information about NicheTrans, please refer to Methods.

### Translating gene expression to metabolites in Parkinson’s Disease (PD)

The Spatial Multimodal Analysis (SMA)^25^ dataset contains brain samples from the Parkinson’s Disease mice, each of which has undergone H&E imaging, Visium sequencing (spatial gene expression measurement), and mass spectrometry imaging (MSI) (spatial metabolic measurement) (Figure. 3A). Additionally, each slice features an intact hemisphere of healthy tissue alongside a lesioned hemisphere modeled to reflect PD through unilateral 6-OHDA administration (Figure. 3A). Here, we used three spatial transcriptomics-metabolomics slices from the originally released datasets (see Methods). Since the two omics were measured on the same slice, spatial metabolomics information can be mapped to the spatial transcriptomics before our analysis (See Methods and Figure. S1B-D). H&E image patches corresponding to the spatial transcriptomics spots were also cropped from the original H&E whole slide images (WSIs). Samples 2 and 3 were utilized to train the model to translate metabolomics from transcriptomics, while Sample 1 was used to evaluate the translation performance (Figure. 3B). With the availability of ground truth (the omics to be translated to), we assessed model performance using metrics including the Pearson Correlation Coefficient (PCC) and Spearman Correlation Coefficient (SPCC) (see Methods). To facilitate a comprehensive comparison, we also tested six state-of-the-art translation methods designed for single-cell multi-omics translation, including totalVI^47^, Polarbear^48^, BABEL^49^, JAMIE^31^, cTP-net^29^, and sciPENN^30^. In terms of our method, we implemented both NicheTrans and NicheTrans*, where NicheTrans utilized the spatial gene expression data, and NicheTrans* additionally incorporated morphology features from H&E image patches. A detailed explanation of the selected single-cell translation methods can be found in Supplementary Notes.

**Figure. 3:**
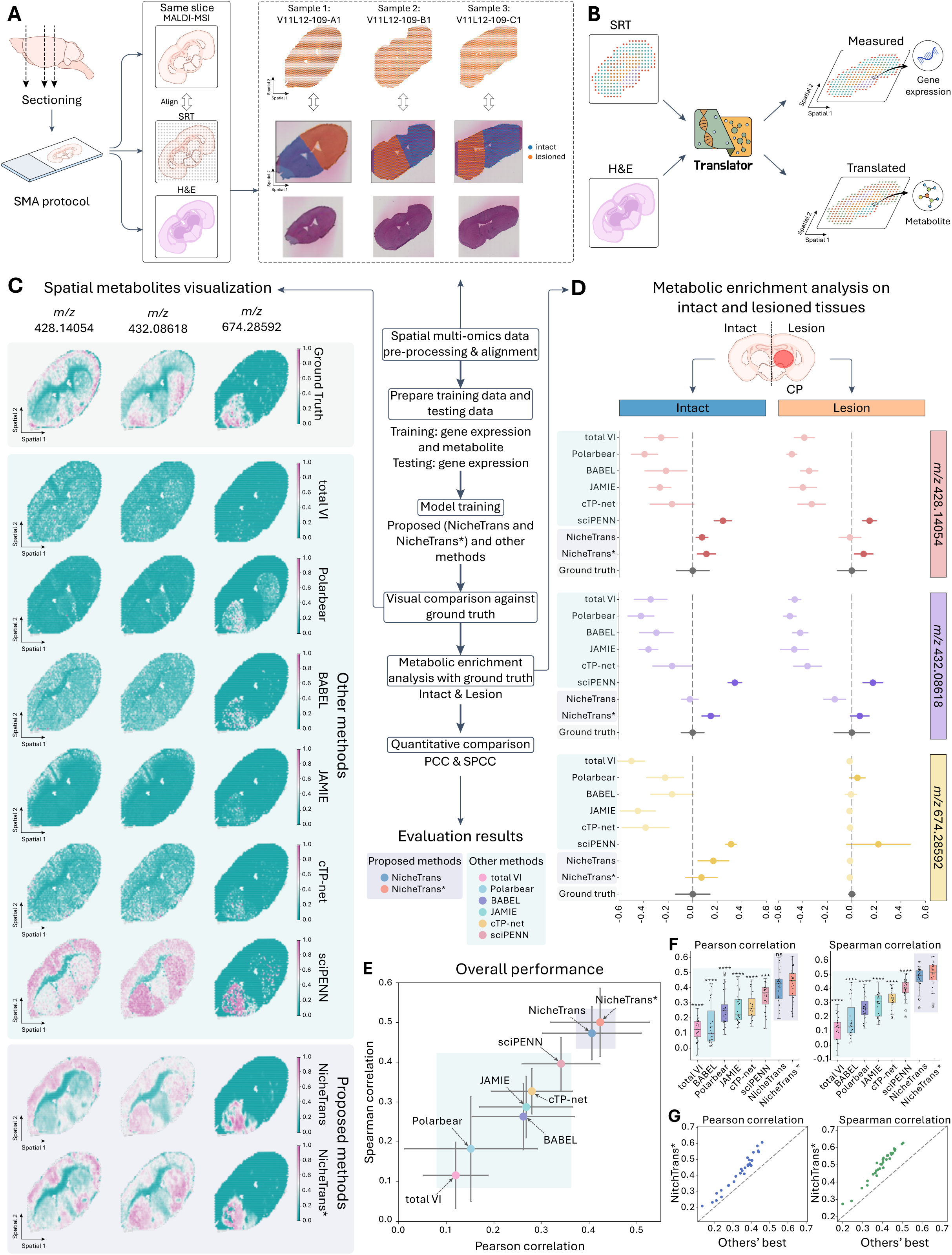
Quantitive and qualititive evalutaion on the SMA dataset. A. Data generation of the SMA dataset. Each slice provides spatial metabolomics data (MALDI-MSI), spatial transcriptomics data, and H&E image data. Spatial metabolomics data were aligned to the corresponding spatial transcriptomics coordinates. B. NicheTrans translates spatial transcriptomics into spatial metabolomics, optionally leveraging the H&E image modality for improved accuracy. C. Visualization of the ground truth and translated results of other methods (single-cell non-spatial translation methods) and our methods (NicheTrans and NicheTrans*) in terms of three selected metabolites. D. Metabolic enrichment analysis on the intact and lesioned tissues, showing mean and standard deviation values to highlight the bias. E. Overall performance comparison, with mean PCC on the x-axis and mean SPCC on the y-axis. F. Performance comparison of different methods in terms of PCC and SPCC using box plots. Each dot in the box plots represents one metabolite. The statistical significance of NicheTrans* and other methods were indicated using * (*: p-value<0.05, **: p-value<0.01, ***: p-value<0.001, ****: p-value<0.0001). G. Comparison of NicheTrans* and the second-best method (sciPENN) on PCC and SPCC.

Qualitative visualization (Figure. 3C), metabolic enrichment analysis (Figure. 3D), and quantitative evaluations (Figure. 3E-G) were performed for benchmarking analysis. In Figure. 3C, we visualized three metabolites that exhibit distinct spatial distributions. The metabolite *m/z* 428.14054 is primarily located in the accumbens nucleus (ACB) and cortex (CTX), the metabolite *m/z* 432.08618 is found in the caudate putamen (CP), and the *m/z* 674.28592 is present in the intact caudate putamen (CP_ intact). The *m/z* 674.28592 has been identified as highly correlative to the PD and its accurate translation is of great significance to a generalized and robust translation model. The first five methods, totalVI, Polarbear, BABEL, JAMIE, and cTP-net, were unable to accurately translate the metabolites from the gene expression data, as indicated by the presence of numerous outliers and significant deviations from the ground truth patterns (Figure. 3C). sciPENN demonstrated improved performance compared to other single-cell translation methods. However, the absence of spatial encoding resulted in worse accuracy than the proposed spatial methods (Figure. 3C). Our proposed NicheTrans and NicheTrans* produced translated results closely matching the ground truth distributions (Figure. 3C). Especially the NicheTrans*, the metabolic expression levels among different functional regions are highly similar to the ground truths (Figure. 3C).

In Figure. 3D, we implemented an enrichment analysis of the selected three metabolites on both intact and lesioned regions (see Methods). We took the ground truth as the center and compared the offsets of different methods. Consistent with the visualization analysis, it is evident that either NicheTrans or NicheTrans* are the closest to the centers compared to other methods (Figure. 3D). All single-cell translation methods except for sciPENN had a relatively lower expression and sciPENN had a much higher expression rate across metabolites, all deviating from the ground truth. This demonstrates the instability and high uncertainty of the single-cell methods in the spatial multi-omics translation process.

In Figure. 3E-G, we quantitatively evaluated NicheTrans, NicheTrans*, and other methods. Figure. 3F utilized the boxplot to display the PCC and SPCC of these methods. Figure. 3G illustrates the one-to-one comparison between NicheTrans* and the second-best method i.e., sciPENN, in terms of PCC and SPCC. Figure. 3E shows the overall performance comparison of different methods using the average values of PCC and SPCC of all the translated metabolites. All these quantitative results highlight the superior performance of NicheTrans and NicheTrans* over the single-cell methods on the spatial omics translations.

### Identifying spatial domains characterized by multi-omics layers

We next demonstrate that the spatial multi-omics data generated by NicheTrans(*) enables the identification of detailed spatial domains that are not detectable using spatial single-omics data alone. Figure. 4A shows the annotated domains of the mouse brain from the SMA dataset^25^. We applied leading spatial domain identification methods (Stagate^50^, GraphST^51^, and SEDR^52^) to analyze both transcriptomic (Figure. 4B) and metabolomic (Figure. 4C) spatial single-omics data separately.

**Figure. 4:**
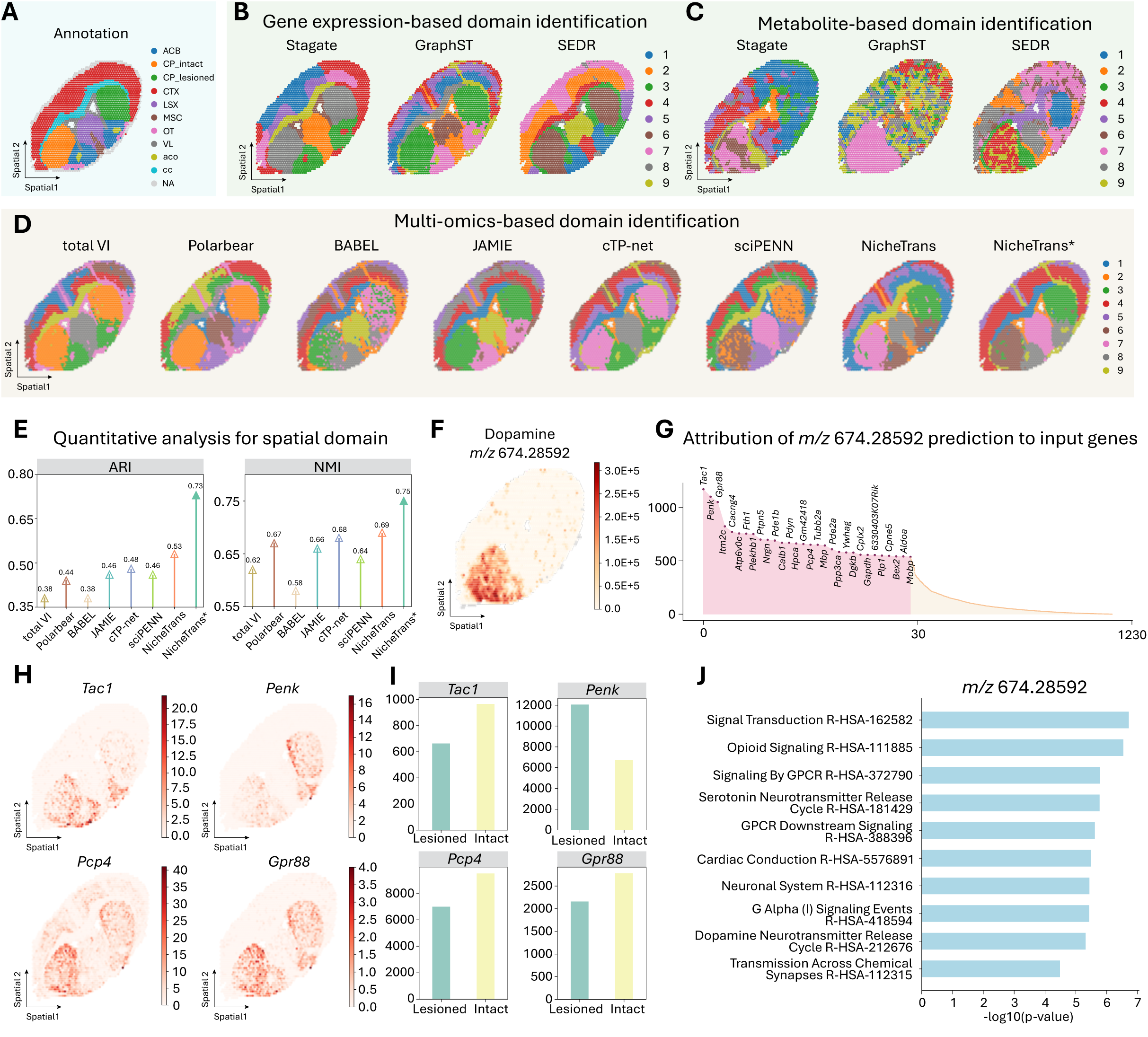
Spatial domain identification and gene programs-metabolite association analysis. A. Spatial domain labels for slice one ‘V11L12-109-A1’. B. Domain identification based on gene expression using state-of-the-art (SOTA) methods. C. Domain identification based on metabolite profiles using SOTA methods. D. Spatial multi-omics domain identification using SpatialGlue, leveraging spatial multi-omics data generated by different methods. E. Quantitative evaluation of the identified domains using ARI and NMI. F. Illustration of the selected PD-related metabolite *m/z* 674.285992 (dopamine). G. Attribution of the *m/z* 674.285992 translation to input gene programs using IG and highlighting the top 30 correlative genes. H. Spatial visualization of the selected PD-related genes, e.g., *Tac1*, *Pcp4*, *Penk*, and *Gpr88*. I. Total expression levels of the selected four genes on both intact and lesioned tissues. J. Gene Ontology (GO) analysis based on *m/z* 674.285992 highly related genes.

Domain identification using transcriptomic data alone fails to distinguish the CP_intact and CP_lesion, regardless of the utilized spatial domain identification methods (Figure. 4B). This could be attributed to the minimal difference in gene expressions of the two brain regions. Although metabolite-based domain identification can marginally differentiate the intact and lesioned hemispheres, the functional regions are poorly displayed (Figure. 4C). These results from Figure. 4B and 4C demonstrate that spatial single-omics data alone cannot fully characterize tissue architectures, highlighting the necessity of spatial multi-omics data for comprehensive biological analysis.

We then evaluated the effectiveness of spatial multi-omics data generated by different methods for downstream analysis. Figure 4D compares domain analysis results using SpatialGlue (an advanced spatial multi-omics analysis method)^27^ applied to data generated by both single-cell (non-spatial) and the proposed spatial translation methods. Among all methods, only NicheTrans and NicheTrans*, combined with SpatialGlue, successfully delineate both functional regions and the two CP variants (Figure. 4D). Quantitative evaluations using the adjusted rand index (ARI) and normalized mutual information (NMI) (two standard metrics used for evaluating spatial domain methods^27,50,53,54^) of the clustering results (Figure. 4E) shows that data generated from NicheTrans* substantially outperforms other methods, reaching 0.73 ARI and 0.75 NMI. These qualitative and quantitative results demonstrate the superior performance of NicheTrans*-generated data for spatial multi-omics analysis, underscoring the critical role of spatial context information in the translation process.

### Developing model interpretation to associate gene programs to the PD-related metabolite

Understanding model interpretation is crucial for elucidating the translation process, particularly the omics-omics relationships in our study. To achieve this, we developed an integrated gradients (IG)-based model interpretation for NicheTrans prediction (see Methods). To demonstrate this, we used a PD-related metabolite *m/z* 674.28592, which shows differential expression between intact and lesioned tissue regions (Figure. 4F), to map its relationships with gene programs.

Figure. 4G represents the top 30 genes identified by the interpretation module. Among the 30 genes, the positively correlated genes *Tac1* (encode a precursor protein correlated to several neuropeptides) and *Pcp4* (encode a calcium-binding protein modulating calcium signaling within neurons), and the negatively correlated gene *Penk* (encode proenkephalin, processed into several neuropeptides), which were particularly highlighted in previous studies^44,55^, were pinpointed. Additionally, other differentially expressed genes *Gpr88*^56^ (encode G protein-coupled receptor 88), *Pdyn* (encode prodynorphin)^57^, and *Hpca* (encode hippocalcin)^58^, which are either explicitly or implicitly correlated to the PD, were also found in this analysis. Figure. 4H and Figure. 4I display the spot expression values in the spatial context and total expression values on the intact and lesioned regions, respectively.

Gene Ontology (GO) analysis of the correlated genes (Figure. 4J) revealed enrichments in key pathways like signal transduction (R-HAS-162582), opioid signaling (R-HAS-111885), signaling by GPCR (R-HAS-372790), serotonin neurotransmitter release cycle (R-HAS-181429), and GPCR downstream signaling (R-HAS-388396). Additional validation was performed on other datasets (Figure. S1A). These findings align with existing research on PD mechanisms. For instance, signal transduction influences PD by mediating oxidative stress, inflammation, mitochondrial dysfunction, autophagy dysregulation, and dopamine signaling disruptions, which collectively contribute to dopaminergic neuron degeneration and disease progression^59^. G protein-coupled receptors (GPCRs) are essential to regulate brain functions by modulating downstream signaling pathways^60^ and have been investigated as novel therapeutics for PD^61^. Together, these interpretation results demonstrate NicheTrans’s ability to identify biologically meaningful relationships between metabolites and gene programs.

### Translating gene expression to proteins in Alzheimer’s Disease (AD)

Co-profiling of protein and gene expression profiles at single-cell resolution has been a great challenge in the spatial omics field. Consequently, this kind of datasets are rarely accessible. In this subsection, we applied the proposed NicheTrans to the spatial multi-omics (gene expression and protein co-profiling) STARmap PLUS dataset^44^. Two brain slices from two AD models and one brain slice from the wild-type model were utilized for extensive analysis (Figure. 5A). For this dataset, we aim to translate the gene expression data to the targeted proteins, i.e., p-tau and Aβ (Figure. 5B), which are the neuropathologic hallmarks of AD^62,63^. Considering the annotated cell types from the original study, NicheTrans provided a flexible approach to incorporate such optional knowledge into the learning process using semantic embedding (Figure. 2) to enhance translation performance (termed NicheTrans*). Our pre-processing steps make omics translation methods flexible to deal with proteins within/outside of cells (see Methods). We trained the model on an AD mouse and tested its performance on another AD mouse and a wild-type mouse.

**Figure. 5:**
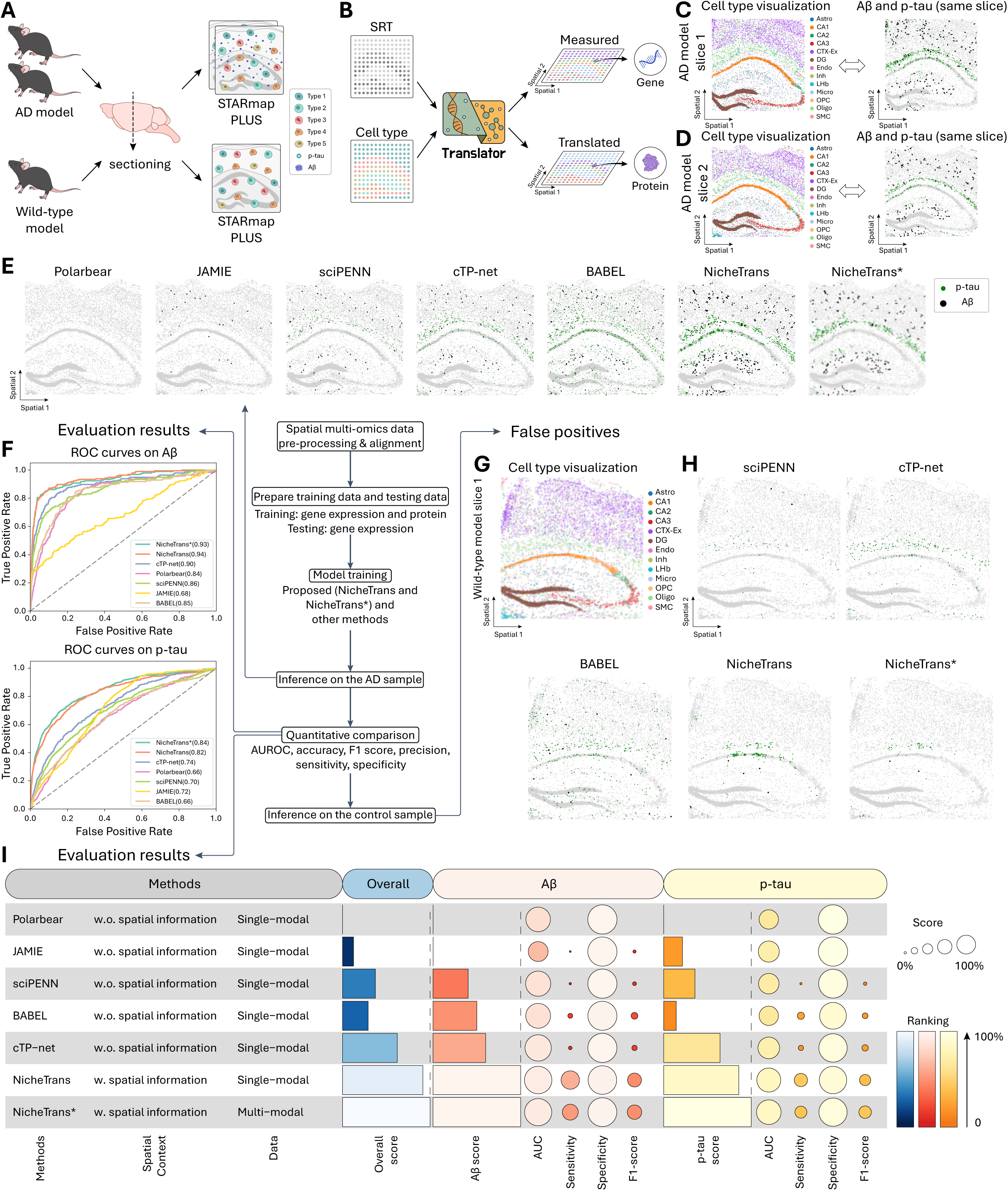
Quantitive and qualititive evalutaion on the STARmap PLUS dataset. A. Data generation of the STARmap PLUS dataset. Each slice is characterized by spatial transcriptomics and spatial proteomics (Aβ and p-tau). There are in total three slices for experiments, including two slices from two AD models and one slice from the wild-type model. B. NicheTrans translates the spatial transcriptomics into spatial proteomics, with the optional incorporation of cell type information for enhanced performance. C, D. Illustrations of AD model slice 1 and AD model slice 2 in terms of cell types and proteins. E. Spatial translation results using different methods on AD model slice 2. F. ROC curves and AUC values for Aβ and p-tau translation across different methods. I. Funky heatmap for Aβ, p-tau, and overall performance evaluation of different methods in terms of accuracy, precision, sensitivity, specificity, F1-score, Aβ score, p-tau score, and overall score (see Methods). G, Illustration of the wild-type model slice 1 labeled with cell types. H, Spatial translation results using different methods on the wild-type model slice.

Figure. 5E illustrates the translation results from five single-cell (non-spatial) translation methods alongside our approaches. For the protein p-tau, both Polarbear and JAMIE show obviously inferior translation performance, identifying only a small number of true positives (Figure. 5E). Although cTP-net and BABEL show improved performance, they still produce a significant number of false negatives, particularly around oligo cells (Figure. S2D). In contrast, the translation results from NicheTrans and NicheTrans* (Figure. 5E) follow highly similar distributions as the ground truth (Figure. 5D). For the protein Aβ in particular, the single-cell translation methods show much lower sensitivity than ours (Figure. 5D-E).

Quantitative evaluations are reported in Figure. 5F and in Figure. 5I. In terms of AUC (Figure. 5F), NicheTrans* outperforms the best single-cell translation method, cTP-net, by 9% and 10% on Aβ and p-tau, respectively (Figure. 5F). Given the large number of cells and the small number of positive ones, different methods show no significant difference in specificity (Figure. 5I). Nevertheless, when it comes to sensitivity and F1-score, NicheTrans and NicheTrans* show much better performance than other single-cell (non-spatial) methods, demonstrating strong identification capability (Figure. 5I). Additionally, we defined the Aβ score, p-tau score, and overall score, the integration of the four metrics (see Methods), to comprehensively evaluate model performance (Figure. 5I). Overall, NicheTrans* demonstrated an improvement over all the methods on the prediction of key proteins of AD (Figure. 5I).

We next rigorously examine whether these methods can avoid false positives in data where target proteins (p-tau and Aβ) are lacking. The testing slice is from a wild-type mouse of the same age (Figure. 5G). Figure. 5H visualizes the translated results from different methods. Polarbear and JAMIE were excluded from the visualization due to their low sensitivities on the AD model (Figure. 5I). Since the wide-type sample is expected to show no p-tau and Aβ distributions, all the positive predictions are false positives. From the results, sciPENN and NicheTrans* demonstrate greater robustness to false positives compared to other models, by translating the fewest false positives (Figure. 5H). When comparing NicheTrans to NicheTrans*, it is evident that incorporating cell type information enhances the model’s performance in minimizing the false positives (Figure. 5H).

### Developing model interpretation to associate cell types and gene programs to Aβ

We next highlight the interpretation capability by attributing the prediction of Aβ to both cell type and gene expression. Consistent with the previous experiments, an IG-based interpretation module (see Methods) was employed for prediction attribution. In Figure. 6C, we visualized the accumulated correlation gradient to different cell types, which highlights the significant correlation between Aβ and Micro, and reveals that Astro and Oligo are the other two important cell types (Figure. 6C). These findings are consistent with the conclusions of the original study^44^ and other studies^63–65^. Moreover, cortical excitatory neuron (CTX-EX) was also thought to be correlated with Aβ. This can be verified by the study that Aβ triggers intermittent aberrant excitatory neuronal activity in the cortex and hippocampus^66^.

**Figure. 6:**
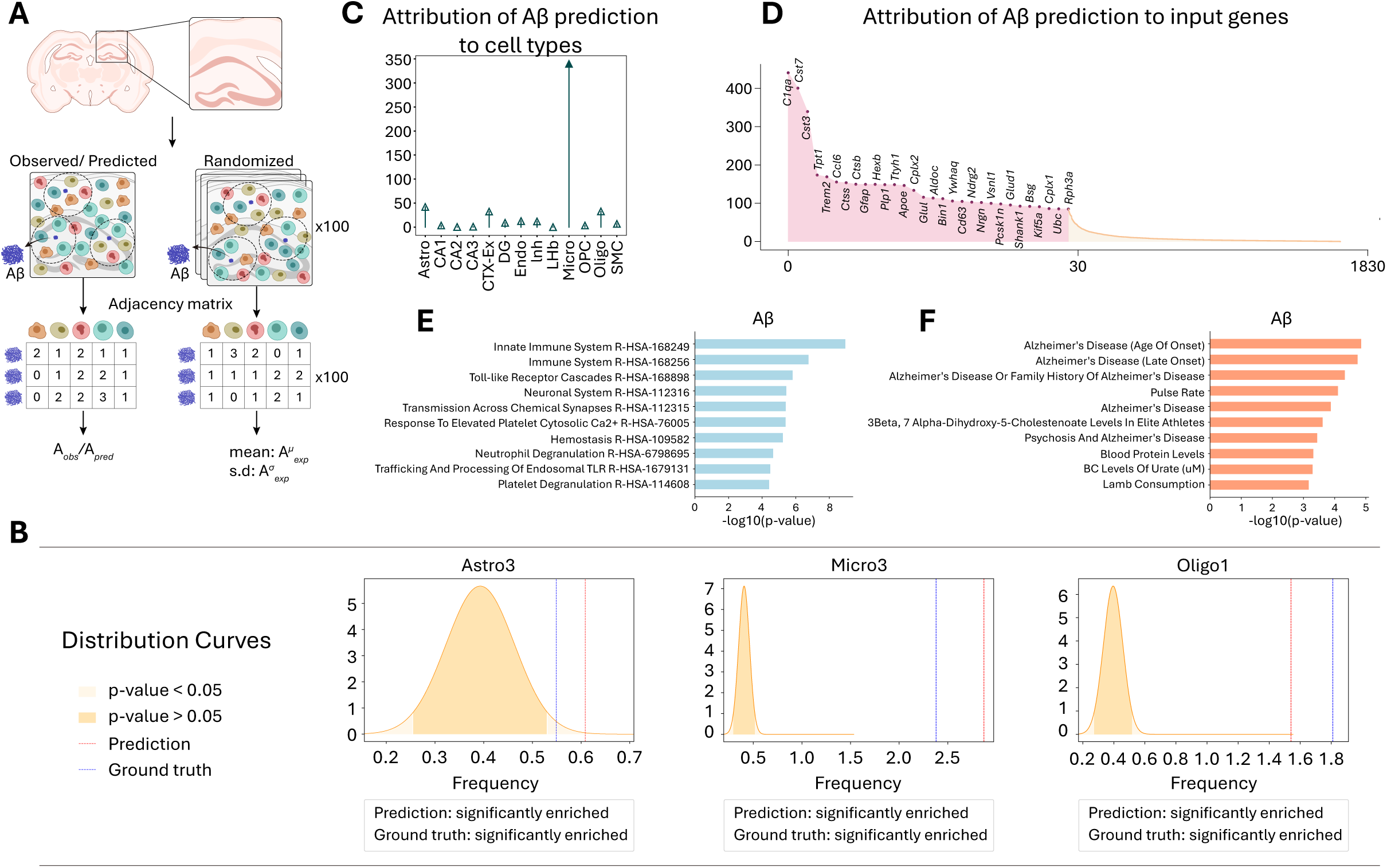
Spatial organization and attribution analysis regarding Aβ. A. Permutation tests quantifying the spatial organization of key cell subtypes associated with Aβ. Adjacency matrices were generated for observed, predicted, and randomized Aβ distributions. B. Comparison of observed and predicted frequencies of different cell types surrounding Aβ, contrasted against a randomized normal distribution. C. Attribution analysis of Aβ prediction to specific cell types. D. Attribution analysis of Aβ prediction to input gene programs, highlighting the top 30 most correlative genes. E. Gene Ontology (GO) enrichment analysis of genes highly associated with Aβ. F. Disease enrichment analysis based on Aβ highly correlated genes.

When attributing Aβ to gene programs, we identified and showed the top 30 correlated genes in Figure. 6D. Zeng et al. particularly emphasize that Micro3 (a subtype of Micro) closely contacts Aβ plaques and is the predominant cell type in correlation to Aβ deposition^44^. Among the 30 genes, all the marker genes of Micro3, i.e., *Cst7* (encode a glycosylated cysteine protease inhibitor related to immune regulation), *Ctsb* (encode cathepsin B related to intracellular proteolysis), *Trem2* (encode the transmembrane triggering receptor expressed on myeloid cells 2 protein), and *Apoe* (encode the apolipoprotein E involved in the metabolism of fats), were all successfully identified. In Figure. S2C, we visualized the top four genes, *C1qa* (encode Complement C1q subcomponent subunit A), *Cst7*, *Cst3* (encode cystatin C protein, that binds Aβ and reduces its aggregation and deposition), and *Tpt1* (encode translationally controlled tumor protein), along with extra-cellular Aβ (grey dots) in the spatial context. It is obvious that *C1qa* and *Cst7* are particularly enriched around Aβ (Figure. S2C), consistent with that the *C1qa*-correlated protein C1q is involved in the pathophysiology of AD^67^.

Furthermore, the GO analysis based on the top 100 identified genes (Figure. 6E) indicates correlations to the innate immune system (R-HAS-168249), immune system (R-HAS-168256), and toll-like receptor cascades (R-HAS-168898). The enrichment analyses on the other datasets are also tested (Figure. S2A). Consistent with previous studies, these pathways are closely related to AD or Aβ. For example, the innate immune system is about the nonspecific part of immunity^68^. Experimental, genetic, and epidemiological data now indicate a crucial role in the activation of the innate immune system as an AD-promoting factor^69^. The broader immune system includes both innate and adaptive immunity that can play different roles in AD development and progression^70^. When implementing the disease association analysis (Figure. 6F), five of the top seven identified diseases are explicitly correlated to AD, significantly validating the reliability of the learned Aβ-gene program correlations. Collectively, these analyses showcase the interpretability of the proposed method.

### Reconstructing the cellular proximities around plaque microenvironment

Beyond the omics-omics and omics-cell type interpretation analysis, we attempted to reconstruct the cellular proximities of the plaque microenvironment using our predicted Aβ. To achieve this, we developed a cell-landmark spatial enrichment analysis based on the permutation test (Figure. 6A). The permutation test is a non-parametric statistical method for testing the significance of observed data distributions by shuffling the data to create distributions under the null hypothesis. This downstream analysis, investigating the spatial organization of the Aβ microenvironment, tells each cell type’s spatial proximity pattern as significantly enriched, significantly depleted, or non-significant.

Specifically, the adjacent matrix between the predicted Aβ plaques and cells was computed based on the spatial coordinates. Using the adjacency matrix (Figure. 6A), we calculated the cell type frequency vector within the spatial neighborhoods of each predicted Aβ plaque. The null distribution is defined by randomly permuting the cell type labels on the whole slice and calculating the distribution statistics. As such, each cell type has an observed frequency and a null distribution around Aβ plaques, which we used to determine the significance (see Methods).

Following the above procedure, we tested each cell type’s spatial proximity pattern towards the predicted Aβ plaques. The results showed that three subtypes—Micro3, Astro3, and Oligo1—are significantly enriched around Aβ plaques (Figure. 6B). These findings agree with the original study that Aβ plaques display a core-shell structure, where Micro3 closely contacts Aβ plaques and Astro3 and Oligo1 are enriched in the outer shells of Micro3^44^. Besides, we also tested each cell type’s spatial proximity pattern towards the measured Aβ plaques, following the same procedure described above. The results using the measured Aβ plaques as spatial landmarks are consistent with those using the predicted Aβ plaques (Figure. 6B). In addition, the reliability of using our predicted Aβ plaques for this analysis was also tested on other cell types (Figure. S3A). These results collectively validate the feasibility of using predicted molecular layer for cell-landmark co-localization analysis, highlighting the utility of our approach in uncovering spatial relationships between cell types and key pathological features.

### Extending to non-brain tissue

Apart from experiments on the highly organized brain, we conducted additional tests on other tissues with complex structures, including breast cancer tissues and human lymph nodes (Figure. 7). Figure. 7A illustrates the data generation process for the breast cancer dataset^35^, i.e., xenium analysis for cell-level spatial transcriptomics and immunofluorescence (IF) staining for spatial proteins. The sequencing and imaging results are shown in Figure. 7B-7C. Towards this spatial multi-omics slice, we took the left part and right part for model training and testing, respectively (Figure. 7B-C). The translation results of CD20 from different methods are illustrated in Figure. 7D. Given the dense distribution of cells, the advantages of NicheTrans over others are not visually obvious. Here, we particularly zoomed in on a selected region to highlight the similarity between ours and ground truth (Figure. 7D). It is evident that the spatial proteomics data translated from NicheTrans exhibits greater spatial continuity and contains fewer over-translated cells. A quantitative comparison of single-cell (non-spatial) methods and ours using PCC and SPCC is shown in Figure. 7E. NicheTrans demonstrates an obvious advantage over others. Additionally, we attributed the translated proteins, i.e., CD20 and HER2, to the input gene programs using the IG-based interpretation module (Figure. S4A). For CD20, its regulation gene *MS4A1* (encode CD20 protein and localize to 11q12) was successfully identified and ranked as the top one important^71^. In terms of HER2, its regulation gene *ERBB2* was also ranked as the top correlative (Figure. S4B). Other genes, such as *CCND1* (overexpressed in HER2-positive cancers and drive cell proliferation), *FOXA1* (correlate with ERα+, GATA3+, and PR+ protein expression as well as endocrine signaling), and *CDH1* (encode Cadherin-1 protein and is a marker gene for breast cancer subtyping), are all also implicitly or explicitly related to HER2 protein and breast cancer. At last, we implemented the trained NicheTrans to its adjacent slice (Xenium replicate 2) (Figure. S4C). We particularly highlighted the expression of CD20 using boxes (Figure. S4C). It is convinced that the translated proteins are highly consistent across slices, demonstrating the strong robustness of NicheTrans.

**Figure. 7:**
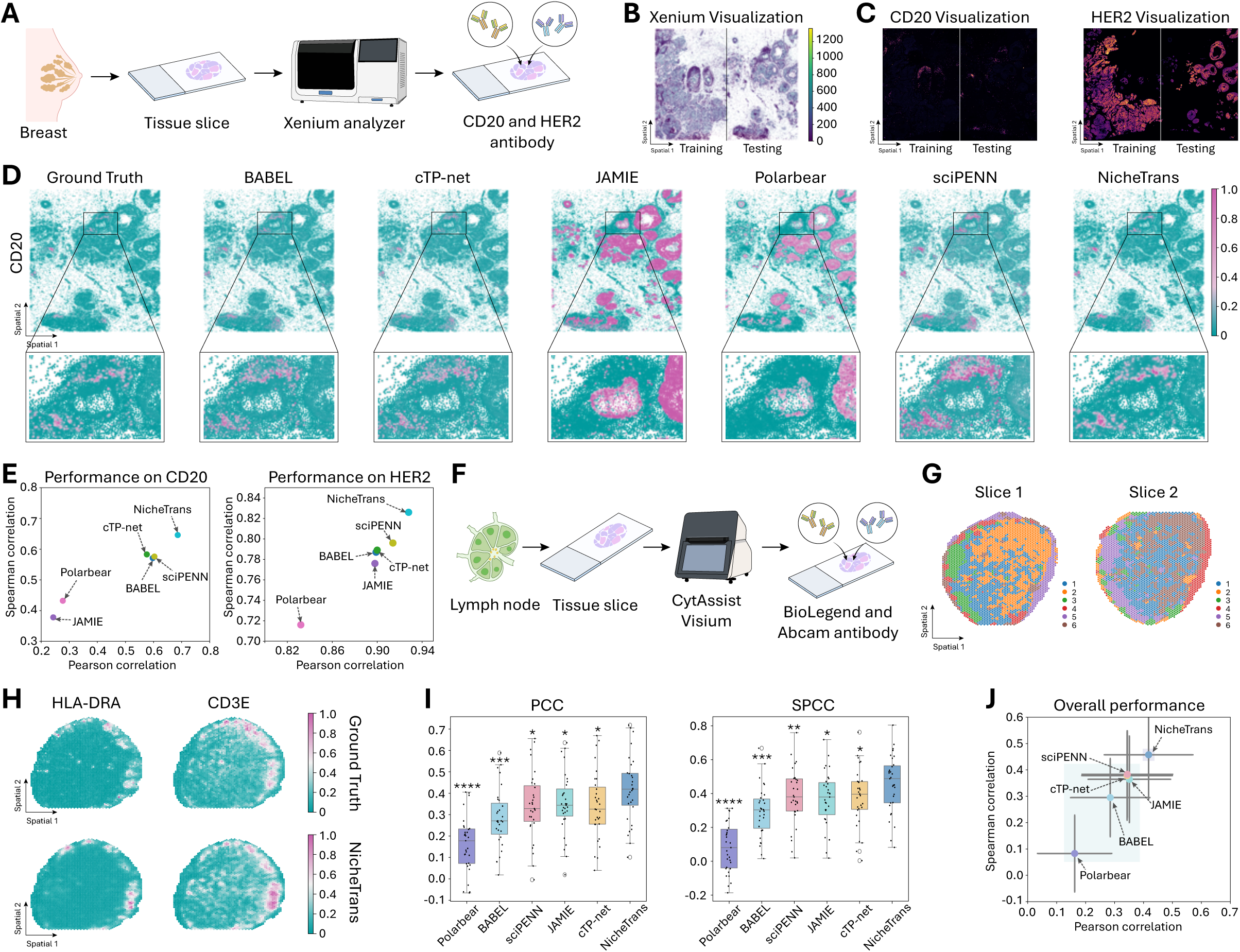
Extending NicheTrans to human breast cancer and human lymph node data. A. Data generation of the breast cancer dataset. Each slice is characterized by spatial transcriptomics using 10x Xenium sequencing and spatial proteomics using immunofluorescence (CD20, HER2, and DAPI antibody). B, C, D. Visualization of the spatial transcriptomics, CD20 protein, HER2 protein in the spatial context. The left part and right part of the slice were utilized for model training and testing, respectively. E. Quantitative evaluation of different methods using PCC and SPCC. F. Data generation of the human lymph node datasets. Each slice is characterized by spatial transcriptomics using CytAssist Visium sequencing and spatial proteomics using immunofluorescence (BioLegend and Abcam antibody). G. Illustration of the two slices with spots labeled by SpatialGlue. H. Spatial visualization of the HLA-DRA and CD3E proteins translated by NicheTrans alongside the ground truth. I. Quantitative evaluation of different methods using PCC and SPCC. The statistical significance of NicheTrans and other methods were indicated using * (*: p-value<0.05, **: p-value<0.01, ***: p-value<0.001, ****: p-value<0.0001). J. Overall performance comparison, with mean PCC on the x-axis and mean SPCC on the y-axis.

Figure. 7F illustrates the data generation process for spatial multi-omics data of the human lymph nodes^27^. The spot-level dataset was processed using CytAssist Visium for spatial transcriptomics sequencing and a 35-plex CytAssist Panel of antibodies (sourced from BioLegend and Abcam) for protein detection (Figure. 7F). There are in total two spatial multi-omics slices used in this experiment, with one for model training and another one for model evaluation (Figure. 7G). In Figure. 7H, we visualized the translated results alongside the ground truth for two proteins (HLA-DRA and CD3E), showing a high degree of translation similarity. The boxplots in Figure. 7I presents a quantitative comparison between different methods using PCC and SPCC, where each point represents a protein. The overall performance comparison in terms of average PCC and average SPCC of proteins is shown in Figure. 7J. Collectively, these results demonstrate the generalization capability of NicheTrans on various tissue types.

## 3. DISCUSSION

In this study, we developed NicheTrans, a Transformer-based multi-modal deep learning framework specifically designed for spatial multi-omics translation. The spatial context information is incorporated into the computation process by formulating each cell and its neighbors together as a niche. Moreover, NicheTrans is highly flexible and scalable, allowing for integrating other data modalities, such as cell type classes and H&E images, as input features to enhance the translation performance. We demonstrated the superiority of NicheTrans for spatial multi-omics translation over other state-of-the-art methods on four benchmarking datasets, highlighting the significance of spatial context information throughout the translation process and the increment brought by the auxiliary multi-modal data.

Using the generated spatial multi-omics data from different methods, we identified spatial domains characterized by integrated gene expression and metabolic signatures that were obscured in single-omics analysis alone. Using the interpretation module, we revealed key relationships between different molecular layers, such as gene programs with dopamine metabolism, gene programs with Aβ, cell types with Aβ, et al. This model interpretation process can help understand the complex biological processes, facilitating the biological experiment designs with more guidance. Furthermore, using the generated spatial multi-omics data, we can quantify the spatial organization of key subtypes, such as glial cells around Aβ in the AD brain model, helping us to understand the biological structures behind various activities.

Although this study has demonstrated NicheTrans’s superior performance in spatial multi-omics translation, several points still need improving. Recently, numerous foundation models have been developed based on large amounts of single-cell omics data^72,73^. These foundation models contain rich knowledge in terms of omics-omics correlation and various biological mechanisms. NicheTrans is expected to be more powerful after incorporating such knowledge or models into the translation process. At the same time, spatial multi-omics sequencing techniques have developed rapidly, enabling more detailed information ranging from omics modalities to text modalities in the future. Incorporating such complex data has not been considered in NicheTrans.

In summary, with the increasing prevalence and significance of spatial multi-omics analysis for various biological processes, NicheTrans represents an economical and efficient method for generating high-quality and informative data for downstream analysis. Additionally, the well-trained neural network can interpret complex correlations between different data modalities, facilitating the in-depth mechanism analysis.

## 4. METHODS

### Data

#### Spatial transcriptomics-metabolomics mouse brain data

The spatial multimodal analysis (SMA) workflow is designed to detect metabolites and transcriptomes in a single tissue section with retained specificity and sensitivity of both modalities (Figure. 3A). The workflow consists of four steps: tissue sectioning, MSI by MALDI, H&E staining and bright field microscopy, and spatial transcriptomics. Unilateral 6-OHDA administration was operated on the brain slices to simulate Parkinson’s disease. This study employed three consecutive slices from one coronal mouse brain for spatial cross-omics translation.

While the SMA workflow enables spatial multi-omics profiling in a single tissue section, the resulting omics data reside on distinct coordinate systems and exhibit differing resolutions (Figure. S1B-C). Consequently, accurate multi-omics alignment is essential before model training and testing. For this dataset, we implemented a two-step alignment strategy: joint landmark labelling followed by a transformation operation. On each slice, eight to ten joint landmarks (indicated by yellow dots in Figure. S1B-C), taking into account relative positions and tissue structure, were manually annotated on both the transcriptomics and metabolomics data. Subsequently, a perspective transformation was then applied to achieve full alignment of the two datasets (Figure. S1D). This procedure was successfully performed on three slices from the ‘V11L12-109’ mouse brain. Detailed visualizations in terms of the pre-aligned and post-aligned omics data are provided in the Figure. S1B-D.

Under the experiment setting of translating transcriptomes to metabolites, two slices were designated for model training and the remaining slice was reserved for model testing. The input transcriptomic panel consisted of the intersection of the top 3,000 highly variable genes across the three slices, and the target metabolite panel comprised the intersection of the top 50 highly variable metabolites shared among them.

#### Spatial transcriptomics-proteomics mouse brain data

STARmap PLUS enables high-resolution spatial transcriptomics along with protein detection in the same tissue section (Figure. 5A). STARmap PLUS sequencing involves five steps: (1) detecting mRNAs and amplifying them as cDNA amplicons; (2) labelling target proteins with specific antibodies; (3) chemically modifying and copolymerizing cDNA, antibodies, and proteins in a hydrogel matrix; (4) reading gene-unique cDNA identifiers; (5) visualizing proteins and their localization. Tissue slices were from two 13-month TauPS2APP triple transgenic mice (an established AD mouse model that exhibits both amyloid plaque and tau pathologies) and a wild-type model as the control (Figure. 5A).

Widespread deposition of extracellular Aβ and intracellular neurofibrillary tangles are the neuropathologic hallmarks of Alzheimer’s disease (AD). Moreover, intracellular tau is released into the extracellular space through processes such as neuronal death and exocytosis. In this case, translating extracellular Aβ and p-tau could be technically challenging. Hereby, we pre-process the STARmap PLUS data by aligning detected proteins with neighborhood cells. Specifically, taking each identified protein p-tau as the center, the nearest three cells within 15 pixels in the image space will be labeled as positive p-tau cells. In terms of the Aβ, the nearest three cells within 50 image pixels will be labeled as positive Aβ cells. The whole labeling process is performed on two slices from the AD models, and the pre-aligned and post-aligned slices are illustrated in Figure. S2B. The intersection of the top 2,000 highly variable genes across the two AD slices were selected as the input gene panel.

#### Spatial transcriptomics-proteomics breast cancer data

The 10x breast cancer dataset comprises two Xenium slices (replicate 1 and replicate 2) with the first slice paired with an IF image (HER2 and CD20 proteins). Given that the spatial transcriptomics data and the IF image residual at different coordinate systems, the original data cannot be directly utilized for the spatial cross-omics translation task. Following the original study, we registered the IF image (replicate 1) to the Xenium slice (replicate 1) through Fiji BigWarp (see the 10x Guide available at https://www.10xgenomics.com/resources/analysis-guides/he-to-xenium-dapi-image-registration-with-fiji). After getting the fully aligned data, we averaged the expressed proteins within each cell as the final cell-level protein expression. The left part and the right part of the replicate 1 were utilized for model training and testing, respectively. All the 313 genes within the designed panel were utilized during the experiments.

### NicheTrans for multi-omics translation

NicheTrans is an innovative Transformer-based multi-modal deep neural network developed for translating between different types of omics data with spatial context, generating spatial multi-omics dataset. NicheTrans is flexible to incorporate additional multi-modal information, e.g., morphological features for H&E images, and cell type classes, to enhance the translation performance from the source omics to the target one (Figure. 2A-B).

The translation process begins by formulating the input slices as niches (the center cell as well as its neighborhoods) based on the spatial coordinates. Each cell within a niche is characterized by its source omics data, which is then encoded into a feature vector using a learnable encoder (the feature encoder in Figure. 2A). This feature encoder comprises two fully connected layers, each followed by one-dimensional batch normalization (*bn*) and a *LeakyReLU* activation function. *LeakyReLU* is particularly advantageous as it introduces non-linearity while mitigating the vanishing gradient problem, enabling the model to learn complex patterns from the data. Subsequently, inspired by the embedding strategy in natural language feature engineering^74^, we applied a semantic embedding operation to integrate the spatial information, cell type classes, and any other relevant prior knowledge of the niches into the computation process (Figure. 2B). Following this, a Transformer block is used to establish long-range dependencies between the central cell and its neighborhoods within each niche, thereby extracting as much information related to the translation process as possible (Figure. 2B). We take the features of the center cell as the niche representation for subsequent computation. At this point, morphological and tissue structure information from H&E images extracted by the image encoder can be integrated into the model, offering a more comprehensive understanding of the tissue’s histological context (Figure. 2A). This histological knowledge can complement the molecular data and help the model better interpret the spatial arrangement and organization of cells. Eventually, the cross-omics decoder, which comprises two linear layers with the first layer followed by a *LeakyReLU* activation function and layer normalization (*ln*), translates the features from each niche into the target omics, e.g., proteins or metabolites.

### Niche construction for spatial multi-omics data

In contrast to single-cell sequencing data, spatial omics data is distinguished by spatial context information, which is crucial for complex in situ analyses. To fully leverage the spatial information during cross-omics translation, each cell and its neighbors within slices are jointly formulated as a niche, allowing the encoding of microenvironment information during the translation process. The niche is defined based on the Euclidean distance. For each cell, its nearest *N* neighbors are identified, with the closest *N*/2 cells classified as first-order neighbors and the remaining *N*/2 cells as second-order neighbors. The hyperparameter *N* was set as 8 for spot-resolution datasets and 12 for cell-resolution datasets.

### Semantic embedding

As previously discussed, semantic embedding is designed for encoding the prior knowledge of niches for feature engineering. In this study, we particularly considered the spatial information and cell type classes (Figure. 2B). In terms of spatial embedding, we introduced three learnable tokens including a central token, a first-order token, and a second-order token. These tokens correspond to the spatial orders defined earlier, where the central cell, first-order neighbors, and second-order neighbors are assigned respective tokens. These tokens were added to the cell features, enriching the spatial contextual information within each niche. For datasets having cell type annotations, we can further conduct cell type embedding. This involved defining a set of cell type-specific learnable tokens, with one token per cell type. For example, in the STARmap PLUS dataset, 13 cell type-specific tokens were defined based on the annotated labels. These tokens were directly added to the cell features, embedding cell type-specific knowledge into the model. Notably, all the tokens were initialized to follow the normal distribution with the mean and std as 0.0, and 0.2, respectively.

### Transformer block

The Transformer architecture has been extensively applied across fields such as medical image analysis^75,76^ and structure biology^77^. In this study, we integrated the Transformer block—a core component of NicheTrans—to model cell-cell communication during the translation process within niches. The Transformer block in NicheTrans is composed of three key components: a linear projection layer, a multi-head self-attention (MHSA) module, and a feed-forward network (FFN), implemented under the pre-layer normalization (Pre-LN) strategy^78^.

As illustrated in Figure. 2B, after the semantic embedding, a linear projection layer first recalibrates the cell features as *F*_1_ = [*f*_1_, *f*_2_, *f*_3_, … , *f*_*N*_] ∈ *R*^*N*×*d*^. Then, the recalibrated features will be fed into the MHSA module. Here, we first generate the query *Q*, key *K*, and value *V*. The generation process can be formulated as follows:

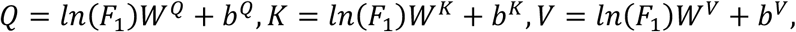

where *W* and *b* represent the learnable transformation matrix and bias, respectively. We keep the multi-head strategy, portioning the features into *m* segments along the feature dimension and getting *Q*, *K*, *V∈R*^*n*×*m*×^(*d*/*m*) . Next, the attention matrix can be generated using the cross-product multiplication between the features and *SoftMax* normalization operation. For simplicity, we set the number of heads as 1 and the whole process can be formulated as follows:

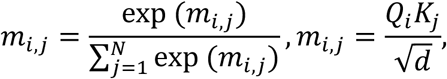

where 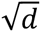 is a scaling factor, and *i* and *j* indicate the indices of *Q* and *K*. Each element in the normalized attention matrix indirectly reflects the communication degree between cells in the omics translation process. The aggregated features are computed as follows:

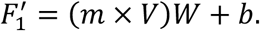

Eventually, the residual between the *F*_1_ and *F*_1_^’^, denoted as *F*_1_^”^, will be taken as the output features.

After the MHSA module, we utilized the FFN for in-depth feature recalibration. Here, the FFN is constructed by two linear layers with the first one followed by the *GEGLU* activation function^79^. Residual operation is conducted between the input features and the recalibrated ones. The feature vector corresponding to the central cell in the niche is selected as the global representation for subsequent decoding computation. The overall computation process of FFN can be formulated as follows:

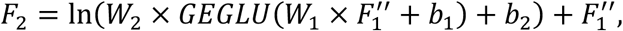

where *W* and *b* indicate learnable matrix and bias, and *ln* is layer normalization.

As emphasized in the Introduction, cellular phenotypes within tissue contexts are shaped not only by intrinsic gene expression but also by interactions with the surrounding microenvironment. The Transformer architecture implemented in NicheTrans effectively models cell-cell communication during the translation process, facilitating a biologically informed and contextually accurate spatial cross-omics translation.

### Image encoder

To extract cellular morphology features from H&E images, we employed convolutional neural networks (CNNs) as the feature extractor. Inspired by the significant success of ResNet^80^ in a wide range of image-based tasks and considering the relatively stable cell morphology features within H&E images, we adopted the lightweight ResNet 18 (18-layer) for feature encoding. Specifically, ResNet18 comprises one convolution block, four residual blocks, and one classification layer. In this study, the final classification layer was removed and the features after the last residual block were regarded as the image representation. Considering the high dimensionality of the extracted image features (512 dimensions), we compress the features into a lower-dimensional representation (128 dimensions) using a nonlinear block which consists of a linear layer, a layer normalization ( *ln* ), and a *LeakyReLU* activation function. The compressed image features were concatenated with the niche representation, jointly as the input to the omics decoder for cross-omics translation.

### Supervised learning for optimizing the neural network

In this study, the proposed NicheTrans was applied to two types of tasks: a regression task for translating the expression levels of target omics and a binary classification task for predicting the expression of target omics. For the regression task, mean square error loss (*L*_*mse*_) is employed to optimize the difference between the predictions and ground truths. The loss function can be formulated as follows:

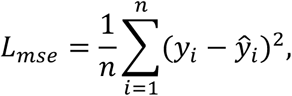

where *y_i_* and 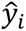 represent the ground truth and predicted expression of the *i*_*t*7_ target omics, respectively, and *n* indicates the length of the target panel. Whereas for the detection dataset, an additional *sigmoid* function is added to the final decoder and the binary cross entropy loss (*L*_*bce*_) was employed for model optimizaation. The loss function can be formulated as follows:

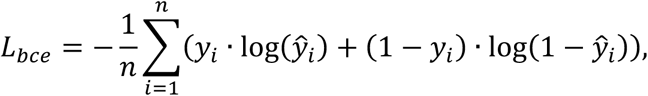

where *y_i_* and 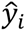 represent the ground truth and predicted probability.

### Evaluation metrics

The regression tasks and classification tasks in this study were evaluated using different metrics. Regarding the first one, Pearson correlation coefficient (PCC) and Spearman correlation coefficient (SPCC) were employed as standards. The PCC can be formulated as follows:

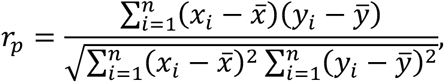

where *x* and *y* represent the translated value and ground truth, respectively, and *x̄* and *ȳ* indicate their mean values. In terms of the SPCC, it can be formulated as follows:

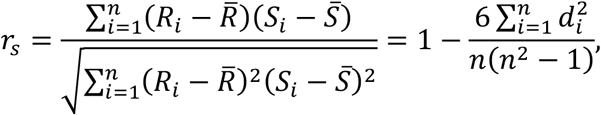

where *d_i_* = *R*[*x_i_*] − *R*[*y_i_*] and *R*[] represents the ranking operation. As for the classification task, the area under the curve (AUC), sensitivity, specificity, and f1_score were employed for model evaluation. These evaluation metrics can be formulated as follows:

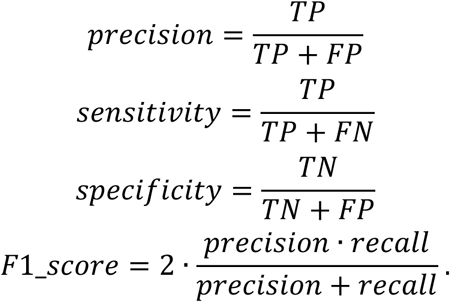

Here, *TN*, *TP*, *FN*, and *FP* indicate true negative, true positive, false negative, and false positive samples, respectively.

In the funky heatmap of Figure. 5I, we utilized the overall score, Aβ score and p-tau score for model evaluation. Referring to the benchmarking study^81^, we sorted values of each evaluation metric in ascending order to get *Rank*_*AUC*_, *Rank*_*sen*_(sensitivity), *Rank*_*spec*_(specificity), and *Rank*_*F*1-*score*_, labelling the best-performing method as N (the total number of methods) and the worst-performing method as 1. Afterwards, the p-tau/ Aβ score can be calculated as:

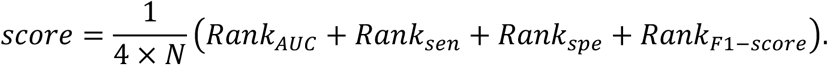

In terms of the overall score, it is the average of the p-tau score and Aβ score. In Figure. 5I, the max-min normalization was processed for the scores to highlight the advantages.

### Datasets for gene expression pattern analysis

To highlight the motivation of this study, we illustrated the cells with identical gene expression patterns on three single-cell resolution spatial transcriptomics datasets undergoing Enhanced ELectric Fluorescence in situ Hybridization (EEL FISH) sequencing^36^, MERSCOPE (Vizgen) sequencing, and Xenium sequencing^35^. Specifically, the EEL FISH slice is from the entire sagittal section of the mouse brain, characterized by the expression patterns of up to 440 genes and 127,591 cells. Regarding the MERSCOPE dataset (https://info.vizgen.com/ffpe-showcase), the selected slice is about ovarian cancer tissue. This slice has in total 251,726 cells with each having 500 genes. At last, the Xenium slice is about the breast cancer tissue^35^, with a total of 167,780 cells each having 313 genes. Before exploring the cells with identical expression patterns, we selected the top 300 highly variable genes (HVGs) using Scanpy to ensure robust analysis and minimize noise.

### Enrichment analysis on the intact and lesioned regions

To facilitate a comparative analysis of the metabolites on both intact and lesion tissues, we conducted the enrichment analysis on the selected metabolites (Figure. 3D). This analysis involves three steps: (1) Max-min normalization was independently applied to each metabolite for both ground truth and the results translated by different methods. This ensured that the metabolite levels were scaled comparably. (2) For the top 100 expressed spots in the intact and lesioned tissues, we calculated the mean and standard deviation for each metabolite across different methods, resulting in statistical values such as *mean*_i*ntact*_g*t*_ , *std*_i*ntact*_g*t*_ , *mean*_*les*i*on*_g*t*_, *std*_*les*i*on*_g*t*_, et al. (3) During the enrichment analysis, the ground truth mean values were subtracted from all the statistical values. This step can highlight the tendency of each method to over- or under-translate metabolite levels, providing insights into their performance relative to the ground truth.

### Model interpretation

For interpreting the well-trained NicheTrans/NicheTrans*, we implemented Integrated Gradients (IG)^82^ for attribution analysis. Supposing we have the input *x* ∈ *R*^*n*^ at hand and baseline *x*′ ∈ *R*^*n*^. IG is defined as the path intergral of the gradients along the straight-line path from the baseline *x*′ to input *x*. Specifically, this process can be formulated as follows:

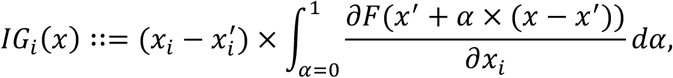

where *F*() represents the neural network. Baseline *x*′ is initialized as a zero-embedding vector by default. Using IG, we can attribute the output to each input element including genes and cell types. In this study, the gradients of all the testing samples were calculated. Given that the positive and negative gradients indicate the positive and negative associations between the outputs and inputs, respectively; here, we sum up the absolute gradients along the sample dimension as the final attribution weights. We utilized IG for transcriptomics-metabolomics analysis on the SMA dataset and for proteomics-transcriptomics and proteomics-cell type analysis on the STARmap PLUS and breast cancer datasets.

### Permutation test

In Figure. 6A, the permutation test was performed to explore the spatial organization of the Aβ microenvironment, highlighting the spatial proximity patterns towards each cell type. Specifically, consider a tissue slice containing a large number of cells, with *m* cells designated as landmarks and a total of *n* distinct cell types. We randomized the cell type of the cells in the slice, resulting in a randomized adjacent matrix *adj*_*ran*_ ∈ *R*^*m*×*n*^. We averaged *adj*_*ran*_ ∈ *R*^*m*×*n*^along the landmark dimension, generating the frequency vector *A_rand_*. Each element in *A*_*rand*_ represents the cell type frequency around the landmark. We proceed with this process one hundred times, summarizing a null distribution with statistical mean 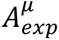 and standard deviation 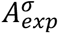. In terms of predicted and measured landmarks, we can also get the frequency vectors *Apred* and *Aobs* based on *adj*_*pred*_ and *adj*_*obs*_, respectively. By comparing *A*_*pred*_ and the null distribution, we can identify the significantly enriched, significantly depleted, and non-significant cell types. By comparing *A*_*pred*_ with *A*_*obs*_, we can showcase the co-localization capability using the predicted landmarks, revealing its spatial organization analysis capability.

### Implementation details

This study was implemented using the PyTorch platform. The models were trained using the Adam optimizer, with the learning rate and weight decay set as 3e-4 and 5e-4, respectively. Model training was conducted for a total of 40 epochs, with a schedule that reduced the learning rate by a factor of 0.1 every 20 epochs. Batch sizes were configured based on dataset resolution: 32 for spot-level datasets and 128 for cell-level datasets. All experiments were performed using a single NVIDIA A800 80G graphics card, ensuring efficient computation for high-resolution spatial multi-omics data.

## Author contributions

Z.Y., J.S., Z.W, S.L, Z.Q., and Y.C. conceived and designed the study, developed the computational methods, performed the analysis, and wrote the manuscript. C.H., Y.L., and J.L. collected and processed the datasets. Y.Z. and R.G. polished the paper writing and designed the figures.

## Acknowledgements

Z.Y. acknowledges the support by National Nature Science Foundation of China (62303119, 32470706), Shanghai Science and Technology Development Funds (23YF1403000), Chenguang Program of Shanghai Education Development Foundation and Shanghai Municipal Education Commission (22CGA02), and Shanghai Science and Technology Commission Program (23JS1410100), National Key R&D Program of China

## Inclusion & Ethics

Not relevant.

